# Real-time motion monitoring improves functional MRI data quality in infants

**DOI:** 10.1101/2021.11.10.468084

**Authors:** Carolina Badke D’Andrea, Jeanette K. Kenley, David F. Montez, Amy E. Mirro, Ryland L. Miller, Eric A. Earl, Jonathan M. Koller, Sooyeon Sung, Essa Yacoub, Jed T. Elison, Damien A. Fair, Nico U.F. Dosenbach, Cynthia Rogers, Christopher D. Smyser, Deanna J. Greene

**Affiliations:** Department of Cognitive Science, University of California San Diego, La Jolla, California 92093, USA; Department of Psychiatry, Washington University School of Medicine, St. Louis, Missouri 63110, USA; Department of Radiology, Washington University School of Medicine, St. Louis, Missouri 63110, USA; Department of Neurology, Washington University School of Medicine, St. Louis, Missouri 63110, USA; Department of Pediatrics, Washington University School of Medicine, St. Louis, Missouri 63110, USA; Data Science and Sharing Team, National Institute of Mental Health, NIH, DHHS, Bethesda, Maryland 20899, USA; Institute of Child Development, College of Education and Human Development, University of Minnesota, Minneapolis, Minnesota 55455, USA; Department of Pediatrics, University of Minnesota Medical School, Minneapolis, Minnesota 55455, USA; Center for Magnetic Resonance Research (CMRR), University of Minnesota, Minneapolis, Minnesota 55455, USA; Masonic Institute for the Developing Brain, University of Minnesota, Minneapolis, Minnesota 55455, USA; Program in Occupational Therapy, Washington University School of Medicine, St. Louis, Missouri 63110, USA; Department of Biomedical Engineering, Washington University in St. Louis, St. Louis, Missouri 63130, USA

**Keywords:** functional MRI, head motion, infant brain, neurodevelopment, neuroimaging

## Abstract

Imaging the infant brain with MRI has improved our understanding of early stages of neurodevelopment. However, head motion during MRI acquisition is detrimental to both functional and structural MRI scan quality. Though infants are commonly scanned while asleep, they commonly exhibit motion during scanning, causing data loss. Our group has shown that providing MRI technicians with real-time motion estimates via Framewise Integrated Real-Time MRI Monitoring (FIRMM) software helps obtain high-quality, low motion fMRI data. By estimating head motion in real time and displaying motion metrics to the MR technician during an fMRI scan, FIRMM can improve scanning efficiency. Hence, we compared average framewise displacement (FD), a proxy for head motion, and the amount of usable fMRI data (FD ≤ 0.2mm) in infants scanned with (n = 407) and without FIRMM (n = 295). Using a mixed-effects model, we found that the addition of FIRMM to current state-of-the-art infant scanning protocols significantly increased the amount of usable fMRI data acquired per infant, demonstrating its value for research and clinical infant neuroimaging.

---

Brain MRI is a powerful tool for studying neurodevelopment during infancy (Graham et al., 2015; Woodward et al., 2006). Head motion during image acquisition is detrimental to both functional and structural MRI, creating a significant hurdle to obtaining high-quality infant MRI data (Cusack et al., 2018; Torres et al., 2020; Zhang et al., 2019). In clinical settings, anesthesia is sometimes used with pediatric populations to reduce head motion during scans. However, concerns about the effects of anesthesia on infants and children (Kamat et al., 2018; Kuehn, 2011; McCann et al., 2019) have motivated both researchers and clinicians to explore alternative motion reduction approaches.

Natural sleep MRI scanning protocols have been developed to help reduce motion in unsedated infants. Feed and swaddle protocols in which the infant is fed immediately before a scan and wrapped snugly in blankets, and use of immobilizers, such as the MedVac Vacuum Splint Infant Immobilizer (Antonov et al., 2017; Golan et al., 2011; Haney et al., 2010; Mathur et al., 2008; Neubauer et al., 2011; Raschle et al., 2012; Weng et al., 2020) have been highly successful. In addition, playing an audio recording of MRI sequence sounds throughout a scan session helps keep infants asleep by preventing changes in ambient noise (Graham et al., 2015; Hughes et al., 2017).

It is generally accepted that the use of these scanning strategies, as well as the participation of a multi-disciplinary and experienced team, is vital for successful image acquisition in unsedated infants (Golan et al., 2011; Graham et al., 2015; Mathur et al., 2008). However, some strategies may not be suitable in all instances, such as feed and swaddle techniques when the risk of respiratory compromise is heightened (Antonov et al., 2017). Further, some infants fail to fall asleep due to environmental factors, such as caregiver anxiety or scanner noise (Antonov et al., 2017; Ellis & Turk-Browne, 2018; Raschle et al., 2012; Tkach et al., 2014). While infant-specific MRI scanners can also increase image quality and decrease motion artifacts (Hughes et al., 2017; Tkach et al., 2014), their acquisition, installation, and maintenance costs are a significant barrier to widespread implementation.

Previously, our group demonstrated that providing MRI technicians with real-time motion estimates via Framewise Integrated Real-Time MRI Monitoring (FIRMM) software reduces the need to overscan, suggesting better efficiency in acquiring high quality, low motion fMRI data (Dosenbach et al., 2017; Fair et al., 2020). In addition, in-scanner head motion can be reduced in children as young as five years old through real-time visual motion feedback provided by FIRMM (Greene et al., 2018). FIRMM calculates and displays a measure of head motion, framewise-displacement (FD), along with other quality metrics, to the MRI technician in real time during a functional MRI (fMRI) scan (Dosenbach et al., 2017). Due to its promise and success in helping acquire low-motion data, FIRMM has been added to many state-of-the-art infant scanning protocols (Fair et al., 2021; Howell et al., 2019). However, the efficacy of real-time motion monitoring for infant MRI scans has not yet been quantified.

In this study, we evaluated the effects of using real-time motion monitoring via FIRMM during acquisition of fMRI data in infants. We measured head motion and the quantity of usable,low-motion fMRI data with and without FIRMM use in infants born prematurely and at term. We hypothesized that real-time motion monitoring would result in the acquisition of more high-quality fMRI data for all infants, as it informs scanner technicians when the infant is holding still well enough to acquire high-quality data.

## Methods

### Participants

Table 1 lists the cohort characteristics for each dataset. Supplementary Tables 1, 2, and 3 report race.

**Table 1.** Participant characteristics for each cohort

| Cohort | Data Collection | Description | Number of infants | Mean gestational age at birth (weeks) | Mean postmenstrual age at scan (weeks) | Female |
| --- | --- | --- | --- | --- | --- | --- |
| 1<br>(term/preterm) | 2007 - 2010 | Healthy term / low grade injury preterm | 83 (9 / 74) | 39 ± 0.6 / 27 ± 2 | 37 ± 0.8 / 38 ± 1.5 | 44% / 58% |
| 2 | 2007 - 2017 | High grade injury preterm | 75 | 25 ± 2 | 39 ± 2 | 39% |
| 3 | 2010 - 2014 | Healthy term | 137 | 38 ± 1 | 38 ± 1 | 56% |
| 4<br>(term/preterm) | 2017 - 2020 | Healthy term / low grade injury preterm | 347 (295 / 52) | 38 ± 1 / 34 ± 2 | 40 ± 2 / 41 ± 1 | 45% / 44% |
| 5 | 2017 – 2019 | Healthy term | 60 | 39 ± 1 | 52 ± 7 | 57% |

Cohort 1 (WUSM): This cohort included 83 infants, with 74 very preterm infants (born at < 30 weeks gestation) prospectively recruited during the first week of life (born at < 30 weeks gestation) and 9 term-born infants (born at ≥ 37 weeks gestation). A total of 84 Infants were recruited from the Barnes-Jewish Hospital Newborn Nursery and Neonatal Intensive Care Unit (NICU) at St Louis Children’s Hospital and scanned between 2007-2010 at term equivalent age (Smyser et al., 2010). One term-born participant was excluded from this study due to brain injury noted incidentally. The preterm infants had no or mild brain injuries, defined as grade I/II using a standardized injury score system (Kidokoro et al., 2013).

Cohort 2 (WUSM): This cohort included 75 very preterm infants (born at < 30 weeks gestation). Infants were recruited from the Neonatal Intensive Care Unit (NICU) at St. Louis Children’s Hospital based upon head ultrasound results obtained in the first month of life and scanned from 2007-2017 at term equivalent age (Smyser et al., 2016). All infants showed evidence of high-grade injury, defined as grade III/IV on a standardized injury scoring system (Kidokoro et al., 2013), and none were excluded from this study.

Cohort 3 (WUSM): This cohort included 137 healthy, term-born infants. A total of 174 infants were recruited from Barnes-Jewish Hospital Newborn Nursery and scanned from 2010-2014 in the first week of life (Smyser et al., 2016). Infants with acidosis on cord blood gas measurements, maternal drug use, and/or incidental brain injury were excluded (n=37).

Cohort 4 (WUSM): This cohort included 347 infants, including 295 healthy term-born and 52 preterm infants (born at < 37 weeks gestation) delivered to mothers prospectively recruited during the second trimester of pregnancy. A total of 382 subjects were recruited from the Barnes-Jewish Hospital Newborn Nursery and scanned while using FIRMM between 2017 and 2020. Term-born infants were scanned in the first weeks of life, while preterm infants were scanned at term equivalent age. A total of 23 infants were excluded due to incidentally noted brain injury of any type and severity, and an additional 12 infants were excluded because FIRMM was not in use.

Cohort 5 (BCP): This cohort included 60 healthy term-born infants from the Baby Connectome Project (BCP) scanned while using FIRMM between 0 and 4 months of age. BCP is an accelerated longitudinal study of children between birth and five years of age, with data collected at the University of Minnesota and the University of North Carolina (Howell et al., 2019). From this dataset, we identified 62 infants scanned within the first four months of life were identified, 2 of which were excluded due to insufficient data collection (less than one completed run of data collected). For subjects that were scanned more than once within the 0 to 4 months age range, the scan session from the earliest time point was included.

### Scanning procedures

Participants from the four WUSM cohorts (1 - 4) underwent a previously established protocol to induce natural sleep during the scan (Mathur et al., 2008). Briefly, a feed and swaddle procedure was used, in which the infant’s feeding schedule was modified to ensure feeding 30 – 45 minutes before the scan time. After feeding, the infant was undressed to a diaper, fitted with ear protection, and snugly swaddled in pre-warmed sheets. The infant was then wrapped in a MedVac Bag, such that when the air was evacuated the infant’s head and neck were held in place. The infant’s head was also stabilized in the head coil with foam pieces. All infants were scanned during natural sleep. Infants were monitored throughout their scan using in-bore cameras, heart rate monitors, and pulse oximeters, as is standard procedure at our institution (Mathur et al., 2008). If an infant woke up during the scan and did not settle, they were removed from the scanner, and repositioned.

Participants from Cohort 5 were scanned during natural sleep following BCP protocols outlined in Howell et al., 2019. Briefly, the infant was fed, swaddled in an MRI-safe blanket, fitted with earplugs and headphones, and rocked to sleep. The infant was then placed on the MRI-safe infant pad on the scanner table and their head was stabilized in the head coil using foam pieces. Infants were directly monitored by a research assistant throughout the scan session. If an infant was unable to fall asleep for a scan, a second scan was attempted. Following this protocol, 1 subject included in this cohort was rescanned.

For Cohorts 4 (WUSM) and 5 (BCP), fMRI data were acquired with FIRMM, which generates real-time motion metrics (framewise displacement [FD]) and displays them to the scanner operator (Dosenbach et al., 2017). The motion data are shown in the form of a FD trace and other quality metrics, such as percent of usable minutes of fMRI data collected below a certain FD threshold (Figure 1). In the time since data collection for this study was conducted, a new version of FIRMM has been developed with similar features (Supplementary Figure 1). For these studies, head motion measurements were computed using an infant specific setting in FIRMM that corrects for a smaller head size radius (r = 35 cm).

**Figure 1.**
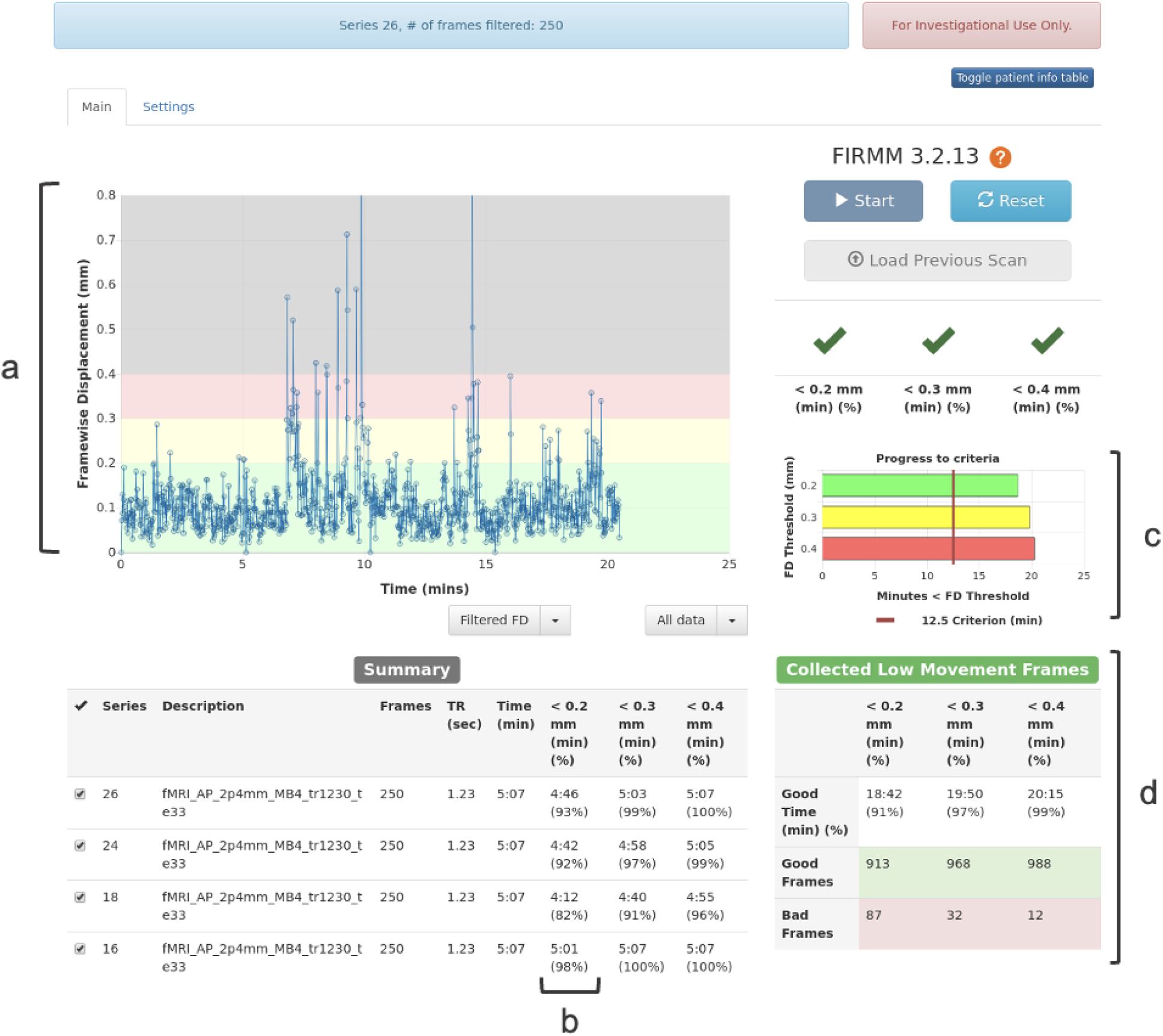
FIRMM Prototype software available in 2017 and used for data collection in Cohorts 4 and 5. (a) *Motion Trace:* plot of FD values for each frame (b) *< 0*.*2 mm (min; %):* minutes and percentage of functional MRI data collected below 0.2 mm for a particular sequence; (c) *Progress to criteria:* total minutes of data collected compared to a predetermined minimum goal of low-motion data; and (d) *Collected Low Movement Frames:* running total of minutes, percent, and number of frames of usable data.

### fMRI acquisitions

For Cohorts 1-4, MRI data were acquired at the Mallinckrodt Institute of Radiology at WUSM or at St. Louis Children’s Hospital (Smyser et al., 2010, 2016). MRI data for Cohorts 1-3 were acquired on a Siemens Trio 3T MRI scanner with a custom 2-channel quad infant head coil manufactured by Advanced Imaging Research Inc. Functional images were acquired using a BOLD contrast-sensitive echo planar sequence; acquisition parameters can be found in Supplementary Table 4. For Cohort 2, rescans were conducted within several days of the initial scan if movement criteria were not met after data processing. Specifically, motion censoring procedures were applied such that frames with FD > 0.25mm or DVARS > 3, in addition to the 2 frames before and after, were removed from analysis, and only contiguous frames of at least 5 were included. Using these criteria, 7 subjects who did not have at least 5 minutes of usable (i.e., low motion) data after one session were rescanned to obtain more data which were analyzed independently.

MRI data for Cohort 4 were acquired on a Siemens Prisma 3T MRI scanner at the Mallinckrodt Institute of Radiology at WUSM with a 64-channel head coil. Functional images were acquired using a BOLD echo planar sequence; acquisition parameters can be found in Supplementary Table 4.

For Cohort 5, MRI data were collected on 3T Siemens Prisma scanners using a 32-channel head coil at the Center of Magnetic Resonance Research (CMRR) at the University of Minnesota and the Biomedical Research Imaging Center (BRIC) at the University of North Carolina at Chapel Hill. Functional images were acquired using the standard Lifespan HCP sequence; acquisition parameters can be found in Supplementary Table 4.

### FIRMM offline processing

In order to generate motion estimates from all cohorts, an offline version of FIRMM was used. FD calculations in the offline and online versions of the software are identical and outlined in detail in Dosenbach *et al*., 2017. In the online version, FIRMM receives DICOMs from the scanner as they are acquired and computes FD by aligning the volumes. In the offline version, FIRMM calculates FD values sequentially using the original BOLD images, prior to any image processing (de-banding and slice-time correction).

### Quantifying data quality

For each participant, we calculated mean FD across all fMRI (BOLD) runs collected. We also measured the amount of low motion (FD ≤ 0.2mm), usable data acquired for each subject across the cohorts to estimate the amount of data that would be retained for analysis in a typical study (Figure 2). We then calculated the percentage of usable minutes of BOLD data per subject (minutes of usable frames / total minutes collected) and used that percentage for further analyses.

**Figure 2.**
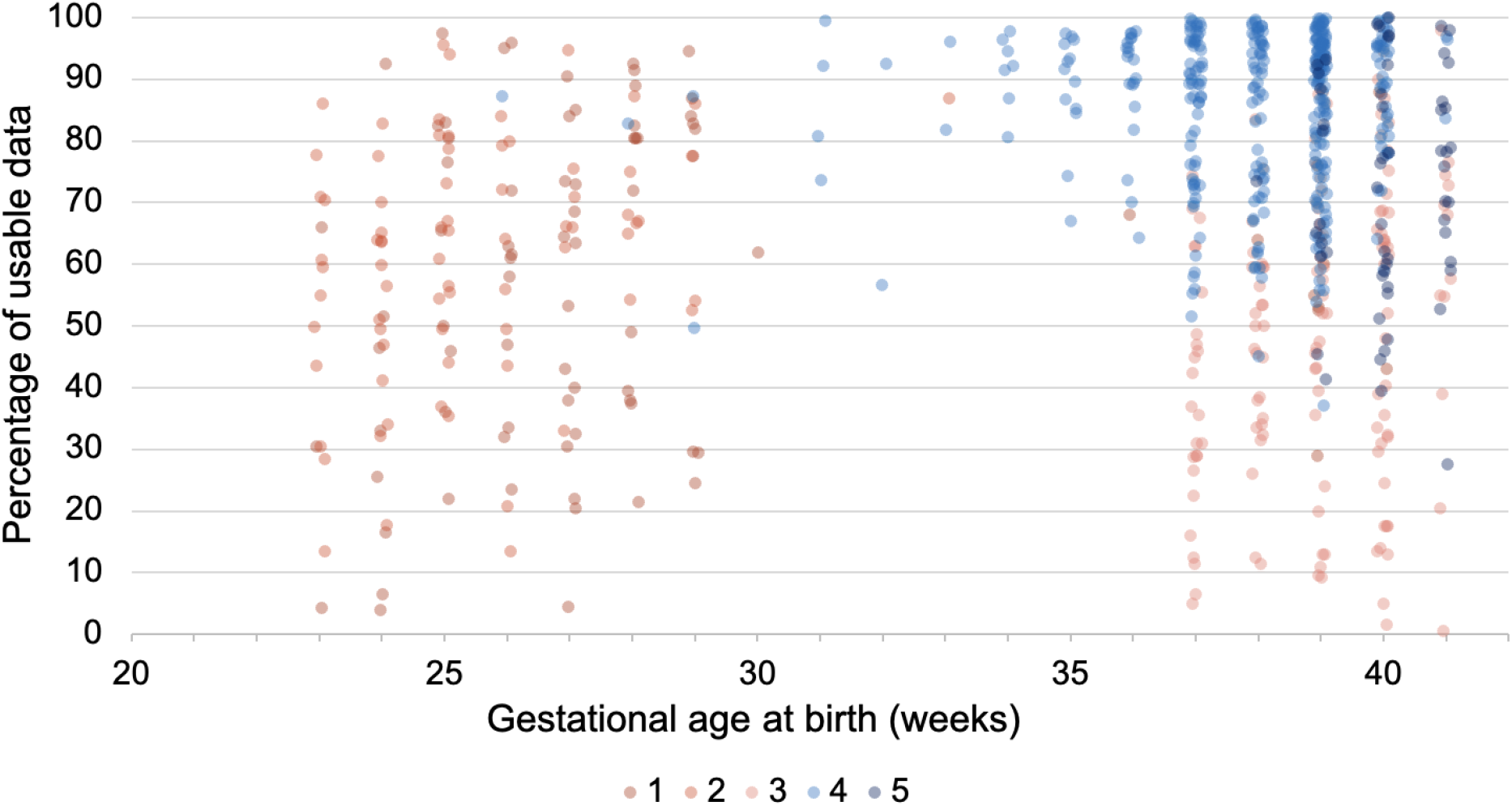
Gestational age and fMRI data quality by cohort. Greater amounts of fMRI data were retained for infants scanned with FIRMM (blue) than for infants scanned without FIRMM (red), independent of gestational age. Data points represent each participant (offset for visualization of overlapping points).

### Mixed effects analyses

Statistical analyses were performed within a mixed-model framework in order to account for potential cohort-specific differences arising for reasons other than the use of FIRMM (e.g., different data acquisition procedures). We modeled each subject’s cohort as a random-effects intercept, while fixed-effects regressors were used to estimate mean FD and percentage of usable data for the cohorts collected with and without the use of FIRMM. Since subject motion is often correlated with age and may also be related to term/preterm birth status, we included gestational age at birth and postmenstrual age at scan as fixed effects covariates. All continuous regressors were mean centered by subtracting the dataset average from the individual regressors prior to inclusion in the model. To test the effect of FIRMM usage on each of the data quality metrics, we included a categorical fixed effects regressor (FIRMM group) indicating whether FIRMM software had been used during data acquisition. Differences in mean FD and percentage of usable data attributable to FIRMM use are denoted Δ FIRMM.

## Results

### FIRMM improves low-motion fMRI data yields in infants

For Cohorts 1-3 (without FIRMM), the mean FD was 0.81 mm and an average of 55% of the data collected from each subject were usable (FD ≤ 0.2 mm; Figure 3). For Cohorts 4 and 5 (with FIRMM), mean FD was 0.26 mm and an average of 79% of the data collected from each subject were usable. Figure 2 and Table 2 report the results for each Cohort. The mixed-effects model revealed a significant reduction in mean FD, t(697) = −10.85, p < 0.001 (Δ FIRMM = −0.52 mm; Table 3a), and increase in percentage of usable data, t(697) = 2.80, p = 0.005 (Δ FIRMM = 21.51%; Table 3b), for cohorts collected with FIRMM (4,5) compared to cohorts collected without FIRMM (1-3), even when controlling for gestational age at birth, post menstrual age at scan, and potential random cohort differences.

**Figure 3.**
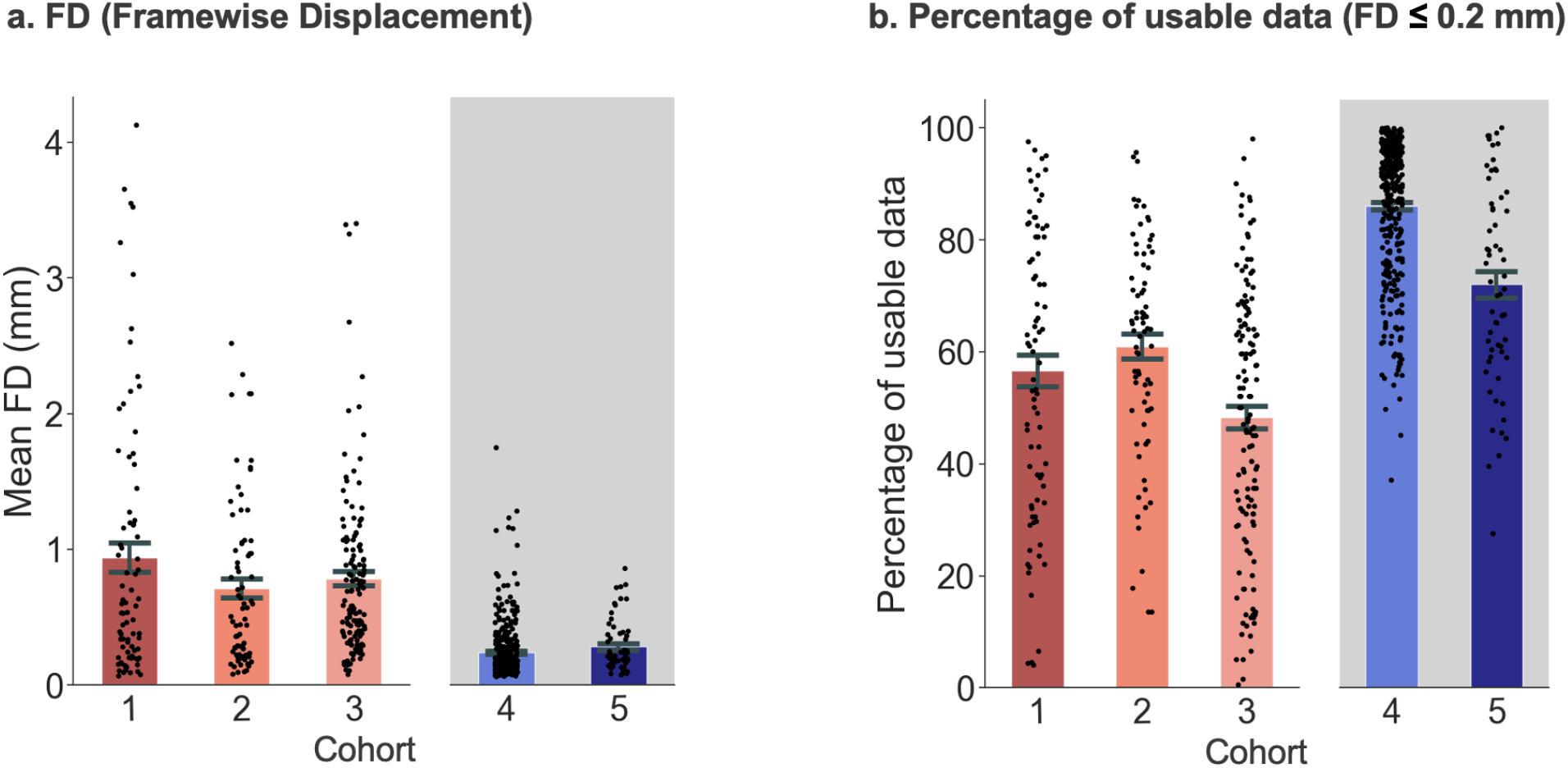
Mean FD and percentage of usable fMRI data collected with (blue) and without (red) FIRMM. (a) Mean FD values; (b) Percentage of usable data defined as frames with FD ≤ 0.2 mm. Each dot (black) represents a subject; error bars indicate standard error of the mean; gray shading denotes cohorts collected using FIRMM.

**Table 2.**
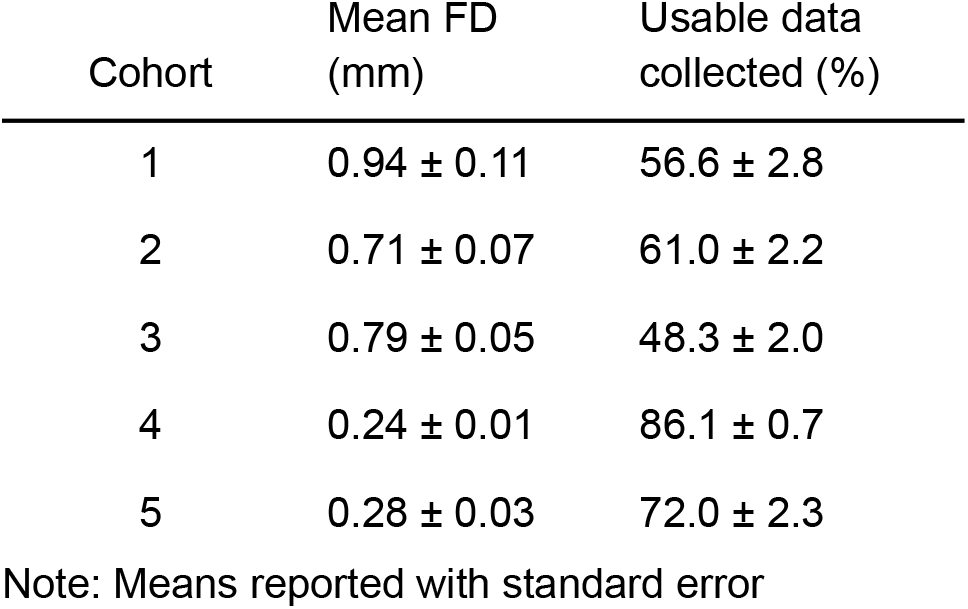
Mean FD and percentage of usable data for each cohort.

**Table 3.** Mixed-effects model results

FD (Framewise Displacement)
| Variable | Estimate (mm) | Standard Error (mm) | tStat | DF | p Value | Lower Bound (mm) | Higher Bound (mm) |
| --- | --- | --- | --- | --- | --- | --- | --- |
| Intercept | 0.79 | 0.033 | 23.33 | 697 | < 0.001 | 0.72 | 0.850 |
| Mean centered GA at birth | -0.01 | 0.004 | -1.84 | 697 | 0.065 | -0.02 | 0.0005 |
| Mean centered PMA at scan | 0.001 | 0.005 | 0.15 | 697 | 0.88 | -0.01 | 0.010 |
| FIRMM group | -0.52 | 0.048 | -10.85 | 697 | < 0.001 | -0.61 | -0.372 |

**b. Percentage of usable data ( $FD \leq 0.2$ mm)**
| Variable | Estimate (%) | Standard Error (%) | tStat | DF | p Value | Lower Bound (%) | Higher Bound (%) |
| --- | --- | --- | --- | --- | --- | --- | --- |
| Intercept | 57.5 | 4.76 | 12.08 | 697 | < 0.001 | 48.16 | 66.85 |
| Mean centered GA at birth | 0.61 | 0.29 | 2.13 | 697 | 0.033 | 0.04 | 1.16 |
| Mean centered PMA at scan | -0.29 | 0.26 | -1.10 | 697 | 0.273 | -0.80 | 0.23 |
| FIRMM group | 21.51 | 7.70 | 2.80 | 697 | 0.005 | 6.39 | 36.63 |

In order to determine whether the apparent FIRMM effect was driven by a single outlier cohort among the three non-FIRMM cohorts, we performed a series of *post hoc* statistical contrasts in which each cohort was compared to the average of the remaining two. Cohort 1 had significantly higher mean FD than the other non-FIRMM cohorts, F(1) = 8.43, p < 0.004 (Table 2), but did not differ significantly in percentage of usable data, F(1) = 0.68, p = 0.4. In order to determine whether or not Cohort 1 was driving the beneficial effect of FIRMM on mean FD, we repeated the mixed-effects analysis for FD while omitting Cohort 1. The difference in mean FD between the non-FIRMM (2 & 3) and FIRMM cohorts (4 & 5) was still significant, t(615) = −13.44, p < 0.001 (Δ FIRMM = −0.52 mm; Supplementary Table 5). Post hoc tests also revealed that the percentage of usable data was significantly lower for Cohort 3 than the other non-FIRMM cohorts, F(1) = 23.02, p < 0.001 (Table 2), but we observed no significant difference in mean FD, F(1) = 0.48, p = 0.49. To determine whether or not Cohort 3 disproportionately contributed to the observed effects of FIRMM on percentage of usable data, we repeated the mixed-effects analysis for percentage of usable data omitting Cohort 3. The difference between the non-FIRMM cohorts (1 & 2) and FIRMM cohorts (4 & 5) remained significant, t(560) = 7.27, p < 0.001 (Δ FIRMM = 25.2%; Supplementary Table 6).

### FIRMM improves fMRI scanning efficiency in both preterm and term infants

The mixed-effects model that included all cohorts also revealed a significant effect of gestational age (Table 3b), suggesting differences between term and preterm infants. Therefore, we tested whether motion and FIRMM effects differed between term and preterm infants. In infants scanned without FIRMM, the preterm infants had a significantly higher percentage of usable data than the term infants, t(292) = 3.65, p < 0.001 (Table 4), but there was no significant difference in mean FD, t(292) = 0.76, p = 0.45. In infants scanned with FIRMM, there were no significant differences between preterm and term infants for either percentage of usable data, t(405) = 1.91, p = 0.06, or mean FD values, t(405)= 0.35, p = 0.73.

**Table 4.** Mean FD and percentage of usable data for preterm and term infants stratified by FIRMM use.

|  | Without FIRMM | With FIRMM |
| --- | --- | --- |
| Preterm | 0.84 ± 0.07 mm<br>58.8 ± 1.9% | 0.26 ± 0.04 mm<br>87.5 ± 1.5% |
| Term | 0.78 ± 0.05 mm<br>48.9 ± 1.9% | 0.25 ± 0.01 mm<br>83.5 ± 0.8% |

Note: Means reported with standard error

To test if there was a relationship between gestational age and FIRMM use, we performed another mixed-effect analysis with the addition of an interaction term for gestational age and FIRMM use. There was no significant interaction for FD, t(696) = 0.21, p = 0.84 (Supplementary Table 7a), or for percentage of usable data, t(696) = −1.62, p = 0.11 (Supplementary Table 7b), suggesting no differential effects of FIRMM use in term and preterm infants.

## Discussion

This study investigated the efficacy of real-time head motion monitoring, using FIRMM software, for improving infant fMRI data quality. We compared metrics of head motion and data retention for five infant cohorts (n = 702), two of which were collected while using FIRMM. We observed that FIRMM use was significantly associated with less head motion (FD: Δ FIRMM = −0.52 mm) and greater data retention (Percent usable data: Δ FIRMM = 21.5%). These results extend previous findings on FIRMM efficacy in children and adults to infants, indicating that monitoring head motion during fMRI data acquisition, in conjunction with other robust scanning procedures, can increase scan quality and efficiency.

Though previous studies have reported successful collection of MRI scans from infants when using infant-specific protocols, the definition of success (or success rate) was qualitative and/or subjective (Torres et al., 2020). For example, studies testing the efficacy of a feed and swaddle procedure on structural MRI scan quality reported 80-95% success rates when defining success as qualitatively providing sufficient information for making a clinical diagnosis (Antonov et al., 2017; Templeton et al., 2020). Another study moved towards a more quantitative approach, using three-point scales for several categories, including overall study quality, presence of motion artifact, spatial resolution, signal-to-noise ratio (SNR), and contrast (Tkach et al., 2014). However, the scales were subjective (e.g., poor, moderate, excellent). In contrast, FIRMM provides objective measures of scan quality for functional brain MRIs. Moreover, one can define success more or less conservatively by selecting a motion (FD) threshold most appropriate for the study goals. Here, we chose a conservative threshold (FD ≤ 0.2mm) that has been shown to mitigate motion artifacts in functional connectivity MRI data (Power et al., 2014).

We demonstrate that real-time motion monitoring provides additive benefit to current gold-standard infant brain scanning protocols. These results raise the question of how real-time motion information was used during the scans to lead to such beneficial effects. Since neonates cannot be coached to hold still in the scanner, study team members (scanner operators, experimenters) actively monitor for increasing movement as an indicator of awakening and then adjust scan protocols accordingly. In fact, the scanner technicians for the WUSM cohorts (1-4) reported using the information from FIRMM in this way, helping them to decide when to intervene with diaper changes or re-swaddling before infants were fully awake and alert. In addition, FIRMM can be used to monitor the quantity of high quality data acquired and make decisions about reacquisitions in order to collect a minimum amount of usable fMRI data, a strategy reported by both the WUSM and BCP study teams. Thus, the benefit derived from FIRMM requires active involvement from the study team during data collection and can involve varying strategies.

Advances in perinatal care have led to increased survival rates as well as improved prognosis for infants born extremely preterm (Hinojosa-Rodríguez et al., 2017). Since many of these infants have diffuse white matter abnormalities (WMA) and subsequent neurodevelopmental irregularities, there is increased interest in obtaining high quality fMRI data to study brain function in this population (Kanel et al., 2021; Ment & Vohr, 2008; Smyser et al., 2010). Many studies have successfully scanned preterm infants for research (Brady et al., 2021; Smyser et al., 2012; Uchitel et al., 2021) and clinical purposes (Ibrahim et al., 2014; Templeton et al., 2020; Woodward et al., 2006). Given that FIRMM use significantly increased the amount of high quality data collected in both preterm and term infants, the addition of FIRMM may be beneficial for studies aimed at advancing our understanding of the neurodevelopmental effects of preterm birth.

In this study, we examined existing datasets in which the entire sample of a given dataset was collected either with or without FIRMM. Therefore, cohorts may have differed in other ways beyond FIRMM use. The particularly detrimental effects of sub-millimeter head motion on functional connectivity MRI were reported in 2012 (Fair et al., 2013; Power et al., 2012; Satterthwaite et al., 2012; Smyser et al., 2010; Van Dijk et al., 2012), leading to additional care and considerations when collecting resting state fMRI data. Given that the non-FIRMM cohorts were collected chronologically prior to the FIRMM cohorts, there were potentially additional updates and improvements in acquisition procedures associated with the new insights about head motion artifacts in the FIRMM cohorts. There was also a difference in age-at-scan in the BCP cohort compared to the WUSM cohorts, such that the BCP cohort was older. To address such potential cohort issues, we controlled for cohort effects in our mixed effects model. We also verified that results were not driven by a single cohort and conducted confirmatory analyses that excluded certain cohorts. These analyses confirmed the significant reduction in head motion and increase in percentage of usable data due to FIRMM, independent of other potential between-cohort factors.

One key benefit of FIRMM is to increase the efficiency of MRI scans. By monitoring head motion in real-time as opposed to computing motion estimates only during post-scan processing, investigators can reduce the need for costly, time consuming rescans, which contribute to sample attrition, by ensuring sufficient data collection. In addition, investigators can collect data until a predetermined criterion is reached (e.g., 10 minutes of low-motion fMRI data). This strategy also reduces superfluous fMRI data collection in low motion participants (over scanning) and allows for the prioritization of other sequence acquisitions (e.g. anatomical, diffusion). For example, real-time motion monitoring has been shown to reduce scan time and associated costs by 57% in individuals with ADHD, Autism Spectrum Disorder (ASD), a family history of alcoholism, and neurotypical controls ages 7-19 years old (Dosenbach et al., 2017). Infant neuroimaging researchers can similarly improve their scan efficiency with the addition of FIRMM to their procedures.

Our results are promising for future neuroimaging studies in infants and may inform research extending the use of real-time motion monitoring to other MR imaging modalities both in research and clinical settings. Most clinical brain imaging in infants implements structural MRI scans, which are also negatively impacted by motion and have more variability in success rates compared to other MRI modalities (Reuter et al., 2015; Torres et al., 2020). Thus, the development of real-time motion monitoring for structural sequences could allow for more efficient use of hospital resources as well as a reduction in the need for anesthesia, helping to avoid the risks and costs associated with sedation. Future research may also benefit from investigating the effects of real-time motion monitoring during awake infant fMRI scanning. The infants in the current study were scanned while asleep, a commonly used approach for minimizing motion and promoting tolerability. However, collecting fMRI data during sleep puts constraints on investigations of active cognitive processes, which requires examining task-evoked activity during wakefulness (Ellis et al., 2020). Moreover, sleep is an inherent confound when comparing infants to children and adults, who almost always undergo research fMRI scans while awake. Therefore, collecting fMRI data from awake infants would allow for the implementation of task fMRI designs, and improve our understanding of neurodevelopment across age. Recent work has demonstrated the feasibility and potential insights that can be gained from awake infant fMRI acquisition (Ellis et al., 2020; Ellis & Turk-Browne, 2018; Yates et al., 2021), and investigators may want to consider using real-time motion monitoring during these scans.

In summary, the addition of real-time head motion monitoring with FIRMM to a gold standard scanning protocol improved the collection of low-motion, highly quality infant fMRI data. The current and previous work suggest that FIRMM improves scanning efficiency by allowing scanner technicians to appropriately intervene when there are substantial motion increases, terminate scans early when sufficient data has been collected, and/or reschedule scan sessions that have a low probability of success. With the potential use of real-time motion monitoring beyond that examined in this study, FIRMM is likely a valuable tool for infant brain MRI scans in both research and clinical settings.

## Acknowledgements

This work was supported by the National Institutes of Health grants R44MH122066 (D.A.F., N.U.F.D.), R44MH121276 (D.A.F., N.U.F.D.), R44MH123567 (D.A.F., N.U.F.D.), K02 NS089852 (C.D.S.), R01 MH113883 (C.D.S. and C.E.R.), R01 MH113570 (C.D.S. and C.E.R.), R01 HD061619, R01 HD057098, P50 HD103525 to the Intellectual and Developmental Disabilities Research Center at Washington University, the Dana Foundation (C.D.S.), the Child Neurology Foundation (C.D.S.), the Cerebral Palsy International Research Foundation (C.D.S.), and the efforts of the UNC/UMN Baby Connectome Project Consortium.

## Contributions

Conceptualization: C.B.D., D.A.F., N.U.F.D., C.R., C.D.S., D.J.G.

Data curation: C.B.D., J.K.K., S.S., E.A.E.

Formal analysis: C.B.D., D.F.M., J.M.K., A.E.M., R.L.M., E.A.E.

Supervision: E.Y., J.T.E., D.A.F., N.U.F.D., C.R., C.D.S., D.J.G.

Writing - Original Draft: C.B.D., D.F.M., D.J.G.

Writing - Review & Editing: All Authors

