## Supplementary Material for "Real-time motion monitoring improves functional MRI data quality in infants"

**Supplementary Table 1.** Self-reported race for Cohorts 1, 2 and 3. Subjects selected one option.

| Race | Cohort 1 (n = 83) | Cohort 2 (n = 75) | Cohort 3 (n = 137) |
|---|---|---|---|
| African American | 39 | 41 | 112 |
| Caucasian | 38 | 33 | 22 |
| Asian | 3 | 1 | 0 |
| American Indian | 0 | 0 | 0 |
| Biracial | 2 | 0 | 3 |
| Other | 0 | 0 | 0 |

**Supplementary Table 2.** Self-reported race for Cohort 4. Subjects were first asked to identify as either "White" or "Other" and then check any of the additional groups that apply.

| Race | Cohort 4 (n = 347) |
|---|---|
| White | 121 |
| Other | 226 |
| Black or African American | 220 |
| Asian Indian | 1 |
| Chinese | 4 |
| Other Pacific Islander | 3 |
| Other | 0 |

**Supplementary Table 3.** Self-reported race for Cohort 5. Subjects were asked to identify their ethnicity as either "Hispanic" or "Non-Hispanic" and identify their race.

| Race | Cohort 5 (n = 60) |
|---|---|
| Hispanic | 6 |
| Non-Hispanic | 54 |
| White | 47 |
| Black | 3 |
| Mixed | 9 |
| Asian | 1 |

**Supplementary Table 4.** Data acquisition parameters

| Cohort | TR / TE (ms) | Flip angle (degrees) | In plane resolution (mm) | Matrix size | Bandwidth (Hz) | Field of View (mm) | Multi-band Factor | Whole brain coverage (slices) | Run length (minutes / frames) | Mean data collected (minutes; range) |
|---|---|---|---|---|---|---|---|---|---|---|
| 1 | 2910 / 28 | 90 | 151 | 64 x 64 | 1662 | x | x | 44 | 4.85 / 100<br>9.7 / 200 | 9.9 (9.7 – 19.3) |
| 2 | 2910 / 28 | 90 | 151 | 64 x 64 | 1662 | x | x | 44 | 9.7 / 200<br>12.125 / 250<br>19.4 / 400 | 16.5 (9.7 – 67.7) |
| 3 | 2910 / 28 | 90 | 151 | 64 x 64 | 1662 | x | x | 44 | 9.7 / 200<br>12.125 / 250 | 10.9 (7.7 – 24.2) |
| 4 | 800 / 37 | 52 | 2 | 104 x 104 | 2990 | 208 | 8 | 72 | 5.6 / 420 | 20.1 (8.3 – 56) |
| 5 | 800 / 37 | 52 | 2 | 104 x 91 | 2990 | 208 | 8 | 72 | 5.75 / 488 | 9.89 (5.6 – 22.4) |

**Supplementary Table 5.** Mixed Effects Model for framewise-displacement (FD), excluding Cohort 1.

| Variable | Estimate (mm) | Standard Error (mm) | tStat | DF | p Value | Lower Bound (mm) | Higher Bound (mm) |
|---|---|---|---|---|---|---|---|
| Intercept | 0.764 | 0.029 | 26.60 | 615 | < 0.001 | 0.708 | 0.821 |
| Mean centered GA at birth | 0.001 | 0.004 | 0.40 | 615 | 0.687 | -0.006 | 0.009 |
| Mean centered PMA at scan | 0.001 | 0.004 | 0.27 | 615 | 0.791 | -0.007 | 0.009 |
| FIRMM group | -0.521 | 0.039 | -13.44 | 615 | < 0.001 | -0.598 | -0.445 |

**Supplementary Table 6.** Mixed Effects Model for percentage of usable data, excluding Cohort 3.

| Variable | Estimate (%) | Standard Error (%) | tStat | DF | p Value | Lower Bound (%) | Higher Bound (%) |
|---|---|---|---|---|---|---|---|
| Intercept | 59.54 | 2.78 | 21.42 | 560 | < 0.001 | 54.08 | 65.00 |
| Mean centered GA at birth | 0.36 | 0.28 | 1.29 | 560 | 0.19753 | -0.19 | 0.90 |
| Mean centered PMA at scan | -0.91 | 0.17 | -5.47 | 560 | < 0.001 | -1.23 | -0.58 |
| FIRMM group | 25.21 | 3.47 | 7.27 | 560 | < 0.001 | 18.40 | 32.02 |

**Supplementary Table 7.** Mixed Effects Model for an interaction between FIRMM use and gestational age (GA) at birth.

**a. FD (Framewise Displacement)**

| Variable | Estimate (mm) | Standard Error (mm) | tStat | DF | p Value | Lower Bound (mm) | Higher Bound (mm) |
|---|---|---|---|---|---|---|---|
| Intercept | 0.784 | 0.03 | 22.45 | 696 | < 0.001 | 0.723 | 0.852 |
| Mean centered GA at birth | -0.008 | 0.004 | -1.82 | 696 | 0.07 | -0.017 | 0.0006 |
| Mean centered PMA at scan | -0.0003 | 0.005 | -0.06 | 696 | 0.95 | -0.010 | 0.012 |
| FIRMM group | -0.5242 | 0.05 | -10.26 | 696 | < 0.001 | -0.621 | -0.424 |
| GA & FIRMM interaction | 0.003 | 0.01 | 0.21 | 696 | 0.84 | -0.021 | 0.031 |

**b. Percentage of usable data (FD ≤ 0.2mm)**

| Variable | Estimate (%) | Standard Error (%) | tStat | DF | p Value | Lower Bound (%) | Higher Bound (%) |
|---|---|---|---|---|---|---|---|
| Intercept | 59.93 | 5.87 | 10.20 | 696 | < 0.001 | 48.40 | 71.46 |
| Mean centered GA at birth | 1.05 | 0.37 | 2.84 | 696 | 0.005 | 0.32 | 1.77 |
| Mean centered PMA at scan | -0.20 | 0.27 | -0.73 | 696 | 0.47 | -0.72 | 0.33 |
| FIRMM group | 20.16 | 9.24 | 2.18 | 696 | 0.03 | 2.02 | 38.3 |
| GA & FIRMM interaction | -0.97 | -0.97 | -1.62 | 696 | 0.11 | -2.15 | 0.21 |

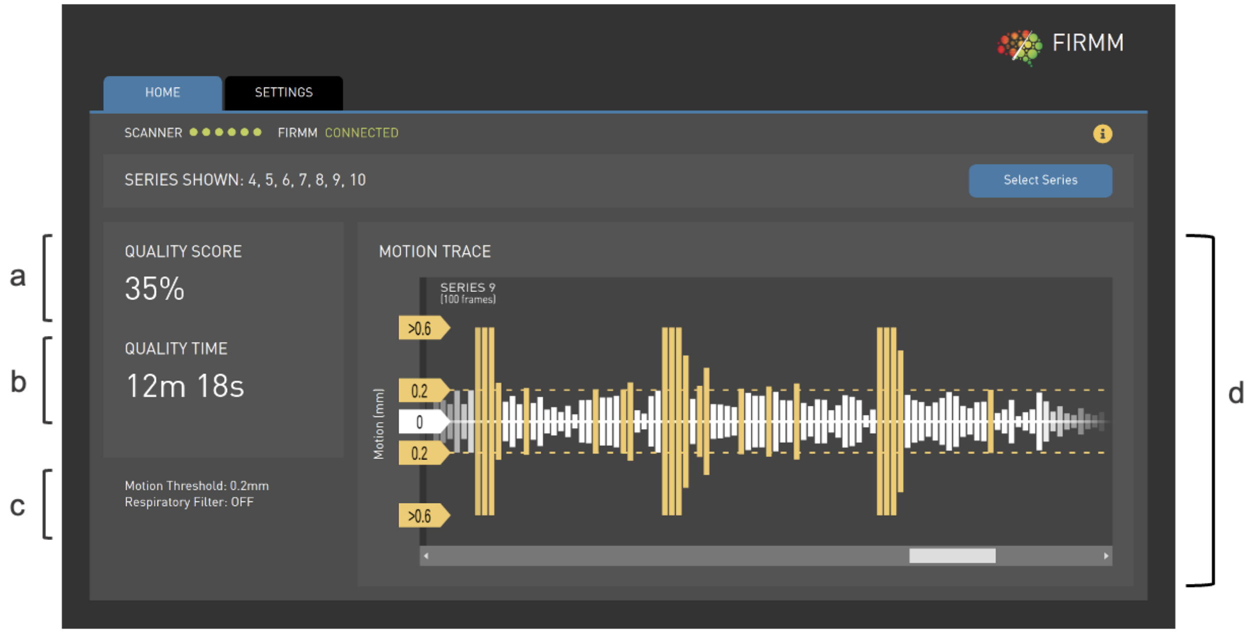


**Supplementary Figure 1.** FIRMM touch-screen interface includes the following features: (a) *Quality Score:* percentage of usable data acquired (b) *Quality Time:* minutes of usable data acquired (c) *Personalized Settings – Motion Threshold:* maximum tolerable FD value (mm) *Respiratory Filter:* filter for artificial motion due to breathing (d) *Motion Trace:* plot of FD values of each frame
